# Evaluating Labelling Efficiency of Commercial SPIONs in Mesenchymal Stem/Stromal Cells for Magnetic Particle Imaging Applications

**DOI:** 10.1101/2025.08.02.668266

**Authors:** Serbay Ozkan, Elena Ureña Horno, Reilteann Saul, Mahon L Maguire, Harish Poptani, Marco Giardiello, Patricia Murray

## Abstract

Magnetic Particle Imaging (MPI) is a state-of-the-art, highly sensitive modality for non-invasive cell tracking. This study evaluated labelling efficiency, biocompatibility, intracellular localization, and MPI detection sensitivity of four commercial superparamagnetic iron oxide nanoparticles (SPIONs)—ProMag, VivoTrax, SynoMag-D, and Ferumoxytol—in mouse mesenchymal stem/stromal cells.

SPION labelling efficiency and cytotoxicity was assessed at varying concentrations and incubation times using Prussian blue staining and ATP-based viability assays, respectively. MPI characterization and transmission electron microscopy (TEM) evaluations were performed for cells labelled for two-hour with ProMag or VivoTrax.

For >90% labelling efficiency, ProMag required 20□µg Fe/mL across all time points. VivoTrax, however, required ≥240□µg Fe/mL, reducing cell viability by >20% necessitating a reduction to 120 µg/mL for further analyses. Transfection agents improved SynoMag-D and Ferumoxytol labelling but compromised viability. MPI analysis revealed linear dependence of signal intensity on labelled cell numbers for ProMag and VivoTrax (r^2^=0.99). ProMag yielded higher signal intensity due to greater iron uptake, although VivoTrax exhibited higher signal per unit iron. TEM confirmed intracellular SPION localization, with ProMag present as individual particles and VivoTrax as aggregates within endocytic vesicles. Low-temperature assays confirmed energy-dependent endocytosis as the primary uptake mechanism. Despite ProMag’s stronger MPI signals and lower detection threshold (12,500 cells), VivoTrax’s superior magnetization per iron suggests its potential following further optimization of cell uptake.

Overall, ProMag and VivoTrax emerged as optimal candidates for MPI-based stem cell tracking. These findings underscore the importance of optimizing both nanoparticle selection and labelling protocols to maximize MPI performance and inform future *in vivo* applications.

## Introduction

Stem cell therapies show promise for treating various diseases due to their regenerative potentials[1]. Among them, mesenchymal stem/stromal cells (MSCs) are particularly advantageous due to their accessibility, multipotency, immunomodulatory capacity, and regenerative secretome[2]. Pre-clinical studies support their therapeutic potential in conditions such as cardiovascular disease, diabetes, renal dysfunction, and autoimmune disorders[1]. However, clinical translation remains limited, with the U.S. Food and Drug Administration (FDA) currently approving MSC use only for graft-versus-host disease[3]. A key barrier is the incomplete understanding of MSC fate post-transplantation, particularly their migration, survival, and integration[4].

Tracking MSC biodistribution is crucial for assessing therapeutic safety and efficacy[5]. Common imaging techniques such as bioluminescence imaging (BLI), nuclear imaging, and magnetic resonance imaging (MRI), each have limitations[6–8]. BLI is sensitive but lacks spatial-resolution and quantitative precision, as signal intensity depends not only on cell number but also on factors like substrate kinetics and tissue depth[9]. Nuclear imaging allows deeper tissue penetration but poses radiotoxicity risks and has a short detection window[10]. MRI with superparamagnetic iron oxide nanoparticles (SPIONs) offers better spatial resolution but suffers from low sensitivity and negative contrast in T2*-weighted images, making it difficult to image SPION-labelled cells in the lungs, liver and spleen[9]. In addition, quantification of the number of SPION labelled cells by MRI is challenging, in part due to the relatively low sensitivity of MRI, but largely due to the fact that SPIONs generate a large volume of negative contrast in T_2_^*^-weighted imaging, obscuring the number of labelled cells within[9]. Magnetic particle imaging (MPI) has emerged as a promising alternative, offering superior sensitivity and quantification without ionizing radiation[11]. MPI signal correlates linearly with iron content, enabling accurate cell quantification[12]. When combined with BLI and anatomical imaging (MRI/CT), MPI enhances real-time tracking of labelled cells[13].

The selection of specific SPIONs as labelling agents is pivotal for the efficiency and reliability of cell tracking[14, 15]. Due to their biocompatibility and strong T_2_^*^ relaxation effects, certain SPIONs, optimized for size, coating, or magnetic properties, are frequently used for MRI-based cell tracking and can also be adapted for MPI signal detection[14–16]. Commercial SPIONs such as ProMag, VivoTrax, SynoMag-D, and Ferrumoxytol vary significantly in these properties[17]. Transfection agents (TAs) like protamine sulphate (PS) can facilitate SPION uptake[12]. Despite their availability, systematic comparisons of SPION labelling efficiency for MPI in MSCs remain limited. Moreover, optimizing the labelling protocol, particularly the incubation period, is crucial to balancing effective SPION internalization with minimizing impact on cell viability. Identifying the shortest incubation time that still ensures efficient labelling is essential to reduce cellular stress and preserve stem cell function, especially in clinical or time-sensitive experimental settings. Therefore, this study investigated the labelling efficiency of the commercial SPIONs and their effects on MSC viability with the aim of identifying suitable candidates for safe and effective MPI-based cell tracking in stem cell research and therapeutic applications.

## Material & Method

### MSC culture

Mouse bone marrow MSCs (D1-ORL-UVA, CRL-12324; ATCC) were cultured in DMEM/F12+GlutaMAX with 10% fetal bovine serum (FBS). Cells at passages 10–15 were used for labelling.

### Cell labelling

The cells were seeded on 12 well plates in growth medium at a density of 1×10^5^ per well and incubated overnight at 37°C in 5% CO_2_. The next day, the medium was removed, and the cells were washed twice with Hanks’ Balanced Salt Solution (HBSS). Separate cohorts of cells were labelled with one of the SPION labelling solutions [ProMag (Bang Laboratories, PMC1N), VivoTrax (Magnetic Insight), SynoMag-D (Micromod, Partikeltechnologie GmbH) and Ferumoxytol (AMAG Pharmaceuticals); physiochemical properties are provided in (Table-1)]. SPION solutions were prepared in a defined concentration range (Table-2), added to each well, and incubated at 37°C in 5% CO_2_ for 18, 4 or 2-hours. For the 18-hour incubation with SynoMag-D and Ferumoxytol, which lack FBS, an equal volume of 20% FBS-containing medium was added to the wells after 4 hours to achieve a standard FBS concentration. TAs [8 IU/mL heparin (5000 IU/mL, Wockhardt) and 240 ug/mL PS (Sigma-Aldrich, P4020)] were added to the labelling media of SynoMag-D and Ferrumoxytol SPIONs.

**Table 1:**
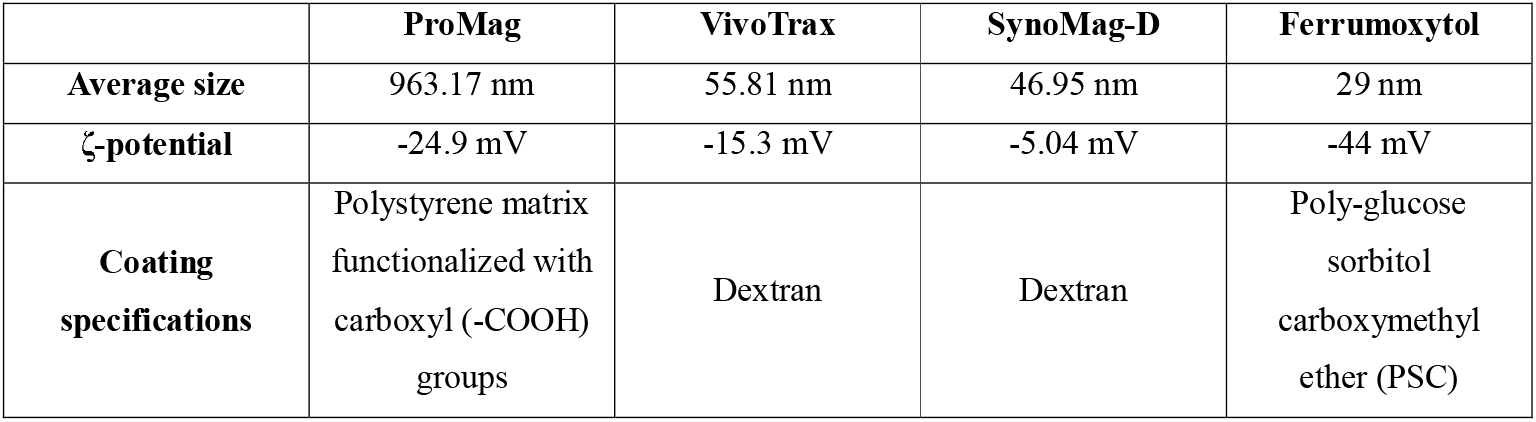
Physicochemical properties of the particles.

**Table 2:** Ranges of iron concentrations in the labelling solutions for the commercial SPIONs.

| VivoTrax in complete medium (10% FBS and 1% Pen/Strep in DMEM/F12) | ProMag in complete medium (10% FBS and 1% Pen/Strep in DMEM/F12) | SynoMag-D in incomplete media (1% Pen/Strep in DMEM/F12 with TAs) | Ferrumoxytol in incomplete media (1% Pen/Strep in DMEM/F12 with TAs) |
| --- | --- | --- | --- |
| Control | Control | Control | Control |
| 30 µg Fe / mL | 20 µg Fe / mL | 67.5 µg Fe / mL | 50 µg Fe / mL |
| 60 µg Fe / mL | 40 µg Fe / mL | 125 µg Fe / mL | 100 µg Fe / mL |
| 120 µg Fe / mL | 80 µg Fe / mL | 250 µg Fe / mL | 200 µg Fe / mL |
| 240 µg Fe / mL | 160 µg Fe / mL | 500 µg Fe / mL | 400 µg Fe / mL |

#### Optimization of incubation periods

To determine the shortest effective incubation time for SPION labelling, preliminary experiments with a single replicate were conducted with cells labelled for 18, 4, and 2 hours. Based on the results of these experiments, two hours was found to offer comparable labelling efficiency across concentrations. No further reductions were deemed necessary. The two-hour condition was then validated in triplicate for statistical analysis.

### Prussian blue staining

Cells were fixed with 4% paraformaldehyde, stained with potassium ferrocyanide and HCl (Sigma Aldrich, HT20) for 30 minutes, and imaged in five random fields at 40× magnification using a Leica DFC420C digital camera. Labelling efficiency was calculated as the percentage of blue-stained cells.

### Cell viability

Cell viability was determined using bioluminescent adenosine triphosphate (ATP) assays. Cells were seeded at 1×10^4^ per well in 100 µL medium and incubated overnight. The next day, media was replaced with SPION labelling solution. Control groups for ProMag and VivoTrax consisted of medium only, without SPIONs. For SynoMag-D and Ferumoxytol, two control conditions were included: medium with TAs but without labelling solution, and medium alone. To detect potential background, cell-free wells were supplemented with defined concentrations of the labelling solutions. After a two-hour incubation at 37°C, the labelling solution was aspirated. After washing, CellTiter-Glo (Promega, G7571) reagent was added, and luminescence measured from triplicate samples with a Fluostar Omega plate reader. Viability was expressed relative to control cells incubated in medium without SPIONs or TAs.

### Magnetic particle imaging

Cells (2.5×10□) were labelled with ProMag (20 µg/mL) or VivoTrax (120 µg/mL) for 2 hours, washed, trypsinized, and counted. Serial dilutions (4×10□ to ∼1×10□) were prepared in 500 µL volumes and imaged individually in a 7-well custom holder using a Momentum^TM^ MPI system (45 kHz, 3 T/m, Magnetic Insight Inc.). Each scan lasted 2.2 minutes. The applied drive field amplitudes were set at 20 mT along the x-axis and 26 mT along the z-axis. Imaging was performed with a field of view (FOV) of 120 mm (Z) × 60 mm (X), and each acquisition lasted 2.2 minutes. Each sample was imaged separately, ensuring no interference from adjacent samples.

### Image analysis

Regions of interest (ROI) were defined using a 5×SD background threshold[18]. Two key sensitivity metrics were analysed: Maximum Intensity and Total Mean Intensity. Maximum Intensity was defined as the highest recorded signal within the dataset, while Total Mean Intensity was calculated by multiplying the mean MPI signal (a.u.) within the ROI by the area of the ROI (mm^2^). Resolution was estimated using the Full Width at Half Maximum (FWHM) (Further detail provided in supplementary method-1). The minimum detectable number of cells was defined as the lowest sample yielding a signal above the established threshold (5×SD of background) following background subtraction of the 2D MPI images.

### Inductively coupled plasma (ICP) iron content analysis

Following MPI, cell pellets were digested in hydrochloric acid (HCl, 36-38%) for 24 hours, diluted with deionized water, and analysed using an Agilent Technologies 5110 ICP-OES system.

### Transmission electron microscopic (TEM) evaluation

Cells were seeded onto a 6-well culture plate and labelled with ProMag or VivoTrax SPIONs as described above. Cells were washed three times, then fixed with 4% paraformaldehyde (PFA) and 2% glutaraldehyde in 0.1□M cacodylate buffer for 1□h at room temperature. After washing, secondary fixation with 1% osmium tetroxide (OsO_4_) was performed for 1□h. Then, the cells were further processed using standard sample preparation procedures, as detailed in supplementary method-2. The samples were examined using TEM (FEI Tecnai G2 Spirit Bio-TWIN) and imaged with a Gatan Rio 16 CMOS camera.

### Low temperature induced endocytosis inhibition assay

To evaluate the cellular internalization mechanism of ProMag and VivoTrax SPIONs, a low-temperature-induced endocytosis inhibition assay was performed[19]. For this, Cells were pre-chilled at 4°C or maintained at 37°C before SPION exposure. After two-hour incubation, uptake efficiency was assessed using Prussian blue staining.

### Statistical analysis

Statistical analysis was performed using SPSS software (version 20.0), and data were presented as mean ± standard error. Differences between groups were assessed using the non-parametric Kruskal-Wallis test followed by the Mann-Whitney U test. For comparison between two groups, a t-test was applied. Significance was defined as p < 0.05

## Results

### Cell labelling and Prussian blue staining

Firstly, the cells were incubated for 18-hour with the defined concentrations of SPIONs, and it was found that 20 µg Fe/mL ProMag, 60 µg Fe/mL VivoTrax, and 125 µg Fe/mL SynoMag-D was able to label more than 90% of the cells, while the highest concentration of Ferumoxytol (400 µg Fe/mL) labelled fewer than 60% of MSCs (Supplementary Figure-1). SynoMag-D and Ferumoxytol were not internalized by the MSCs in the absence of TAs. To determine whether effective labelling could be achieved more in a shorter time period, the incubation time was reduced to 4 hours. This resulted in higher minimum SPION concentrations being required to exceed 90% labelling efficiency—240□µg/mL for VivoTrax and 500□µg/mL for Synomag-D—except for ProMag, which maintained >90% efficiency even at the lowest tested concentration. Ferumoxytol peaked at only 33% efficiency at 200□µg/mL (Supplementary Figure-2). Further shortening the incubation to 2 hours, ProMag achieved 100% efficiency at 20□µg/mL, while VivoTrax, Synomag-D, and Ferumoxytol reached 80%, 77%, and 46% at 240, 500, and 200□µg/mL, respectively (Figure□1).

**Figure 1:**
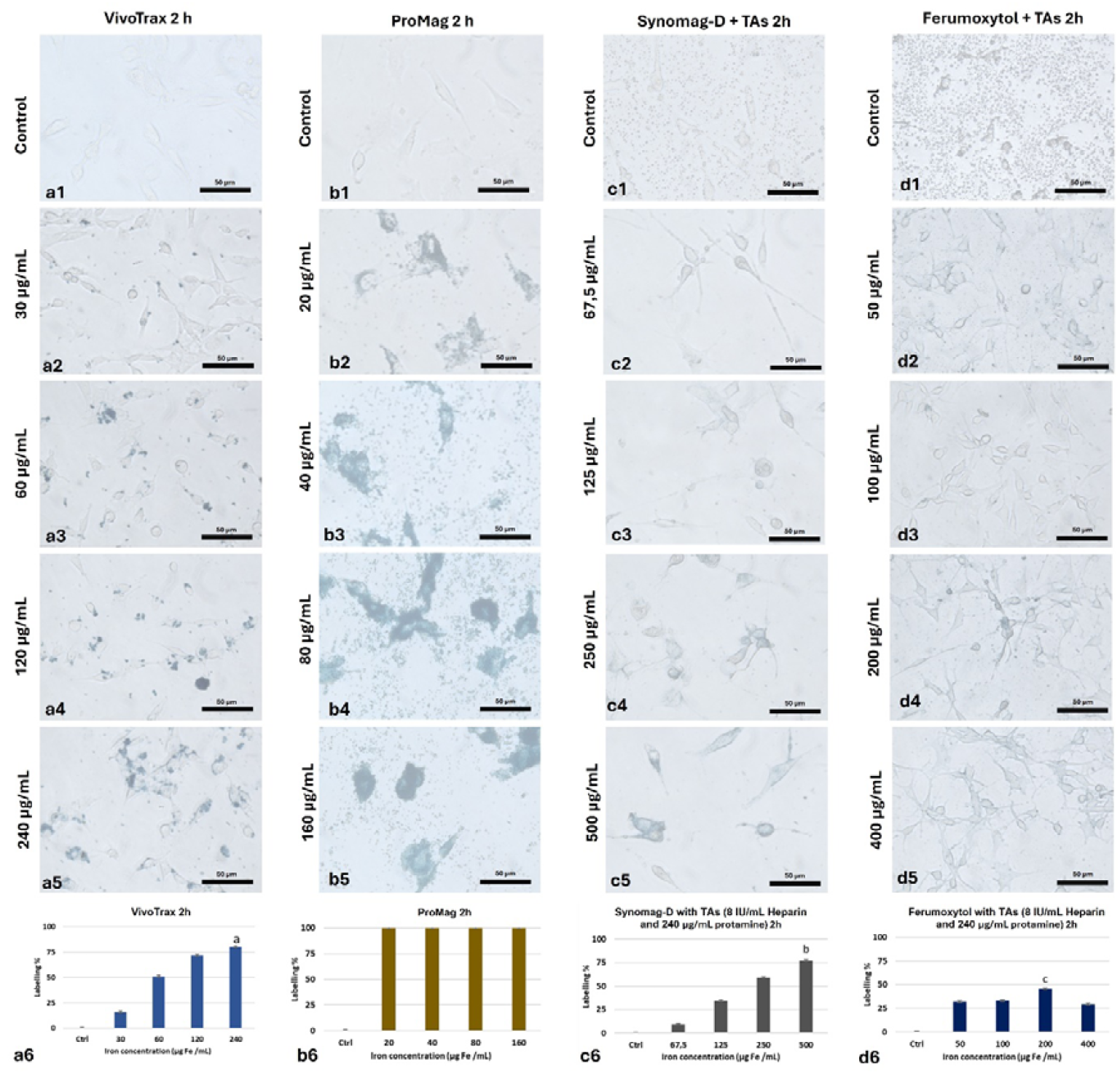
(a-d) Representative images of adherent MSCs incubated with commercial SPIONs at different concentrations, and graphical expression of labelling efficiencies (a6, b6, c6, and d6). *P*^*a*^ < 0.05 vs Ctrl and 30 µg Fe /mL; *P*^*b*^ < 0.05 vs Ctrl, 67.5-and 125-µg Fe /mL; *P*^*c*^ < 0.05 vs Ctrl.

### Cell viability

ATP-based assays showed >90% viability up to 120□µg Fe/mL VivoTrax, with a drop to 60% at 240□µg/mL (Figure□2a). ProMag showed dose-dependent toxicity, with highest viability (84%) at 20□µg/mL (Figure□2b). For SynoMag-D, viability peaked at 125□µg/mL and dropped at 500□µg/mL; for Ferumoxytol, highest viability occurred at 400□µg/mL, while the lowest was in the control group (including only transfection agents) (Figures□2c–d).

**Figure 2:**
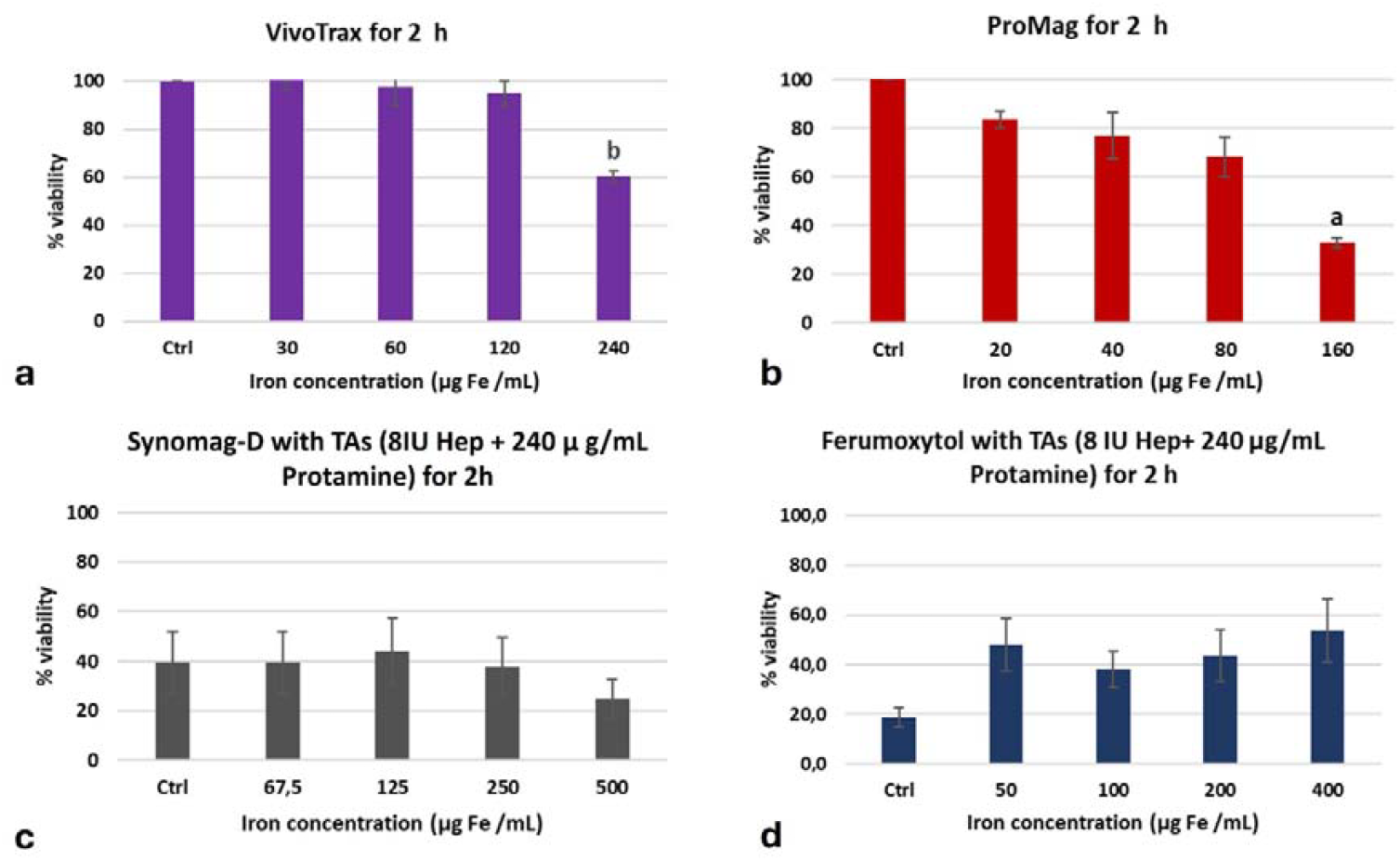
(a-d) Graphical representation of ATP-based viability assay. *P*^*a*^ < 0.05 vs Ctrl, and 20 µg Fe/mL; *P*^*b*^ < 0.05 vs Ctrl.

### MPI and ICP iron content analysis

Due to low viability and uptake with SynoMag-D and Ferumoxytol, only ProMag (20□µg/mL) and VivoTrax (120□µg/mL) were used in subsequent experiments. Prussian Blue staining confirmed labelling efficiencies of 97% (ProMag) and 83% (VivoTrax) at a two-hour incubation period (Supplementary Figure□3). Labelled cells were distributed into Eppendorf tubes in decreasing numbers by two-fold dilutions, and 2D MPI projection images were captured (Figure 3). Minimum detectable cell numbers were 12,500 for ProMag and 50,000 for VivoTrax (Figure□3). At the highest tested cell number (4□×□10□), iron content was 11.2□µg (28.00□pg/cell) for ProMag and 2.3□µg (5.75□pg/cell) for VivoTrax. For both ProMag and VivoTrax, the maximum MPI signal (r^2^=0.99 and r^2^=0.98, respectively) and total mean signal intensities (r^2^=0.99 for both) were linearly correlated with the number of labelled cells (Figure□4a-b). ProMag-labelled cells exhibited higher maximum and mean signal intensities than VivoTrax (Figure□4a–b). However, VivoTrax produced a stronger MPI signal per unit of internalized iron (Figure□4c–d), indicating that, despite lower overall iron uptake, each iron particle in VivoTrax-labelled cells contributed more efficiently to signal generation compared to ProMag. MPI resolution, measured by FWHM, was slightly better for VivoTrax (9.88□mm) than ProMag (10.81□mm) (Figure□4e).

**Figure 3:**
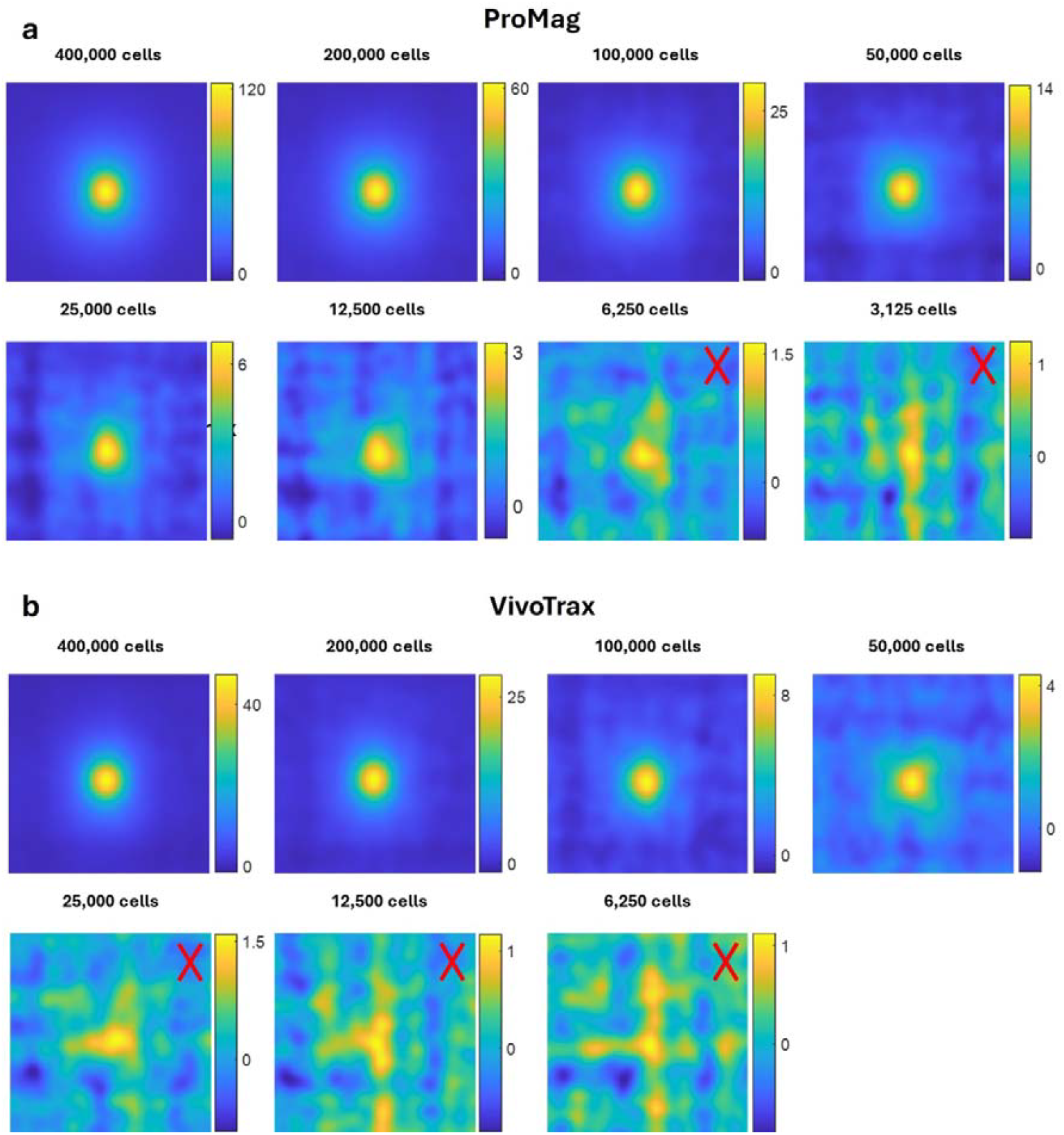
(a-b) 2D MPI images of cell labelled with ProMag (a) and VivoTrax (b) following subtraction of background. An ‘X’ marks images where the signal was not distinguishable from background.

**Figure 4:**
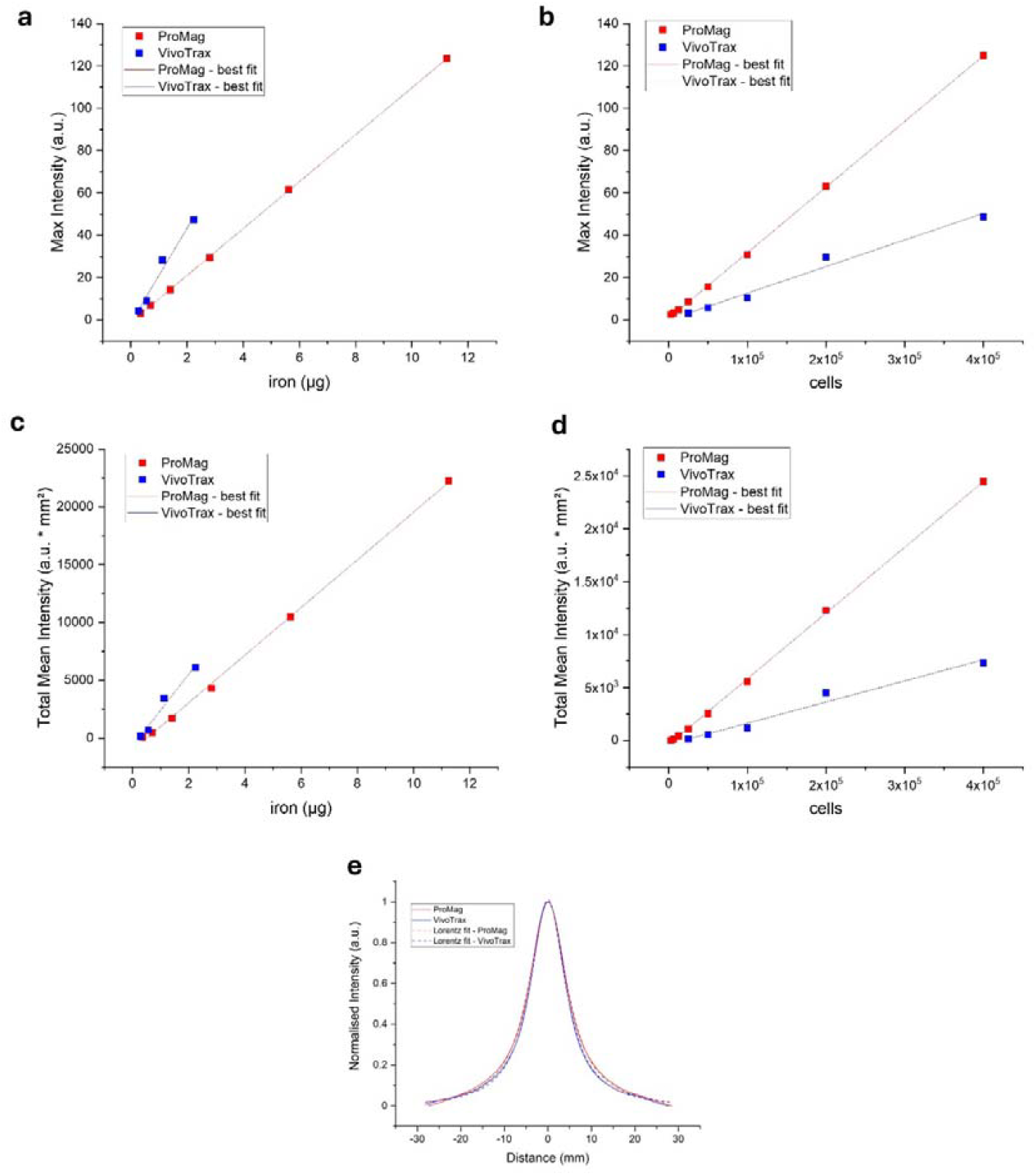
(a-e) MPI signal as a function of cell number and iron mass for ProMag (in red) and VivoTrax (in blue): (a) Maximum intensity and (b) total mean Intensity as a function of cell number, and (c) maximum intensity and (d) total mean intensity as a function of iron content, (e) fitted Lorentzian curve fitting for resolution of labelled cells with either ProMag or VivoTrax.

### Low temperature induced endocytosis inhibition assay

Cellular uptake of both ProMag and VivoTrax was significantly reduced by 81.5% and 69.3%, respectively, under low-temperature conditions relative to 37 °C (Figure-5), confirming active internalization rather than mere surface adherence.

**Figure 5:**
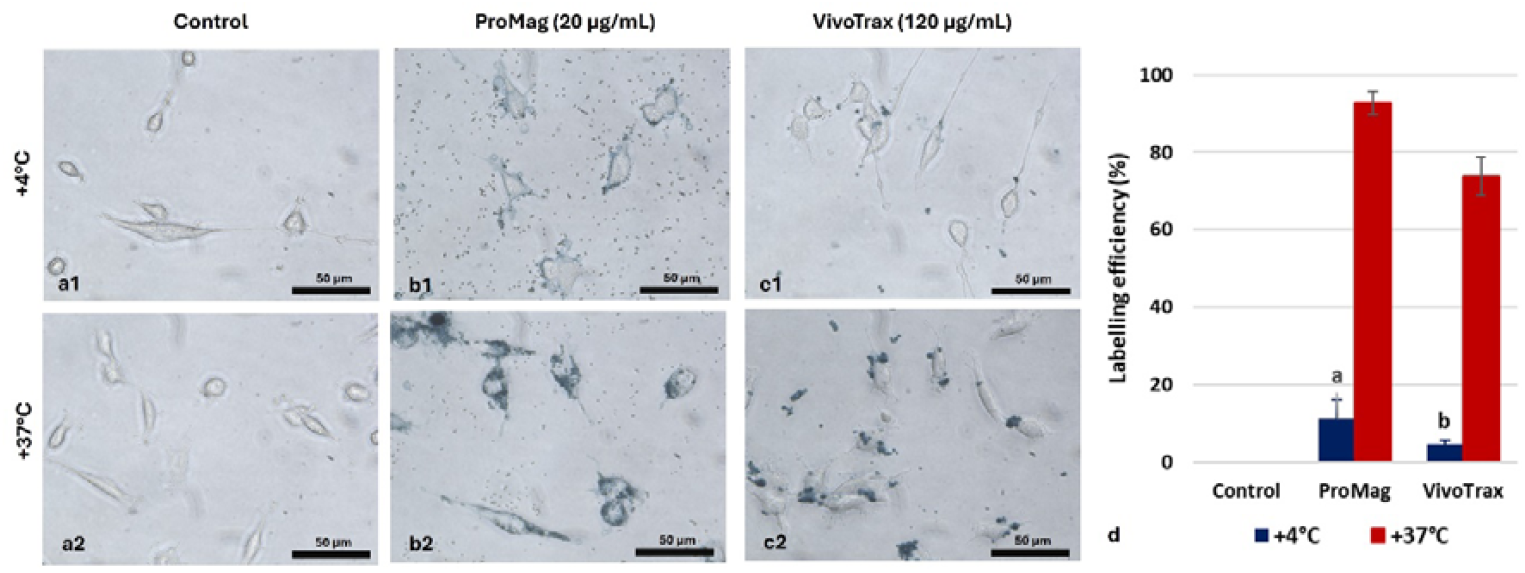
(a-c) Low temperature induced endocytosis inhibition assay. Representative figures for Prussian blue staining of MSCs incubated at +4°C and +37°C in the absence of labelling particle (a) and in the presence of ProMag (b) or VivoTrax (c). (d) Graphical expression of labelling efficiencies. *P*^*a*^ < 0.001 vs ProMag +37°C and *P*^*b*^ < 0.001 vs VivoTrax +37°C.

### TEM evaluation

TEM evaluations of VivoTrax labelled cells showed aggregated particles were inside the endocytotic vesicles. Free particles and membrane-attached particles were also observed, the latter potentially representing the initial stage of the endocytic process (Figure 6a). VivoTrax SPIONs were also detected in the cytoplasm, either freely dispersed or localized within various cellular compartments such as mitochondria and the smooth endoplasmic reticulum. TEM evaluation of ProMag particles showed that they were substantially bigger than the VivoTrax particles. While not necessarily aggregated, they were often found in close proximity to each other and were predominantly observed in the cytoplasm, typically opposite the nuclear region (Figure-6b).

**Figure 6:**
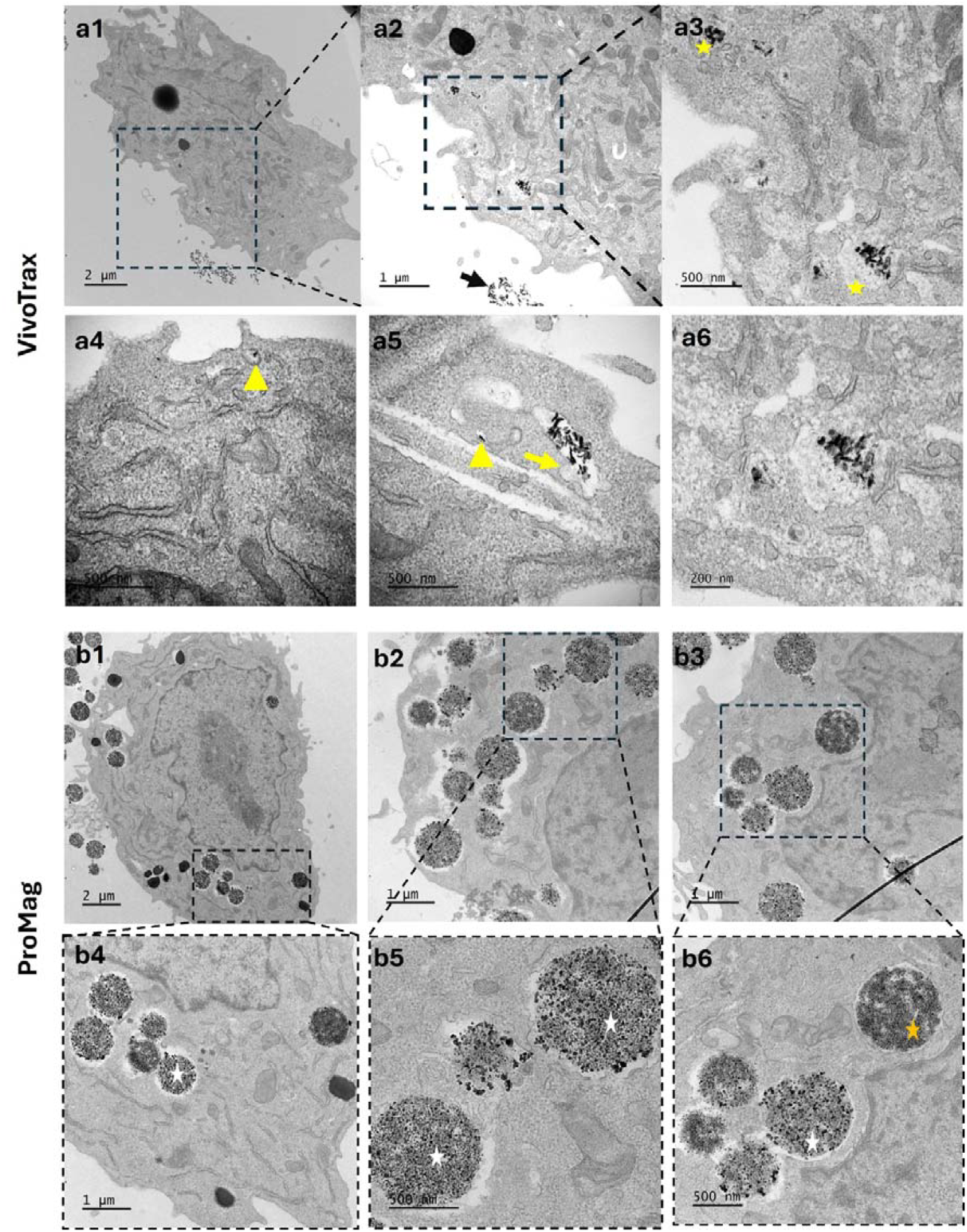
(a-b) Transmission electron microscopic evaluation of VivoTrax (a) and ProMag (b) labelled mMSCs. Yellow star: late endosomes including VivoTrax SPIONs; black arrow: unbound SPIONs; yellow arrow: early endosome fusing with vesicles containing SPIONs; yellow arrowhead: a particle encapsulated in vesicle (early endosome); white star: endocytosed ProMag particles; orange star: ProMag with granulated appearance.

## Discussion

In this study, we evaluated four commercially available SPIONs—ProMag, VivoTrax, SynoMag-D, and Ferumoxytol—for labelling mMSCs for MPI. ProMag and VivoTrax emerged as the most promising candidates based on labelling efficiency and cell viability. ProMag, specifically, produced a higher MPI signal per cell, suggesting superior sensitivity for detecting small cell populations.

Since prolonged SPION exposure can impair MSC viability and function, short incubation times are preferable, particularly for clinical applications requiring rapid and scalable protocols. Initial single-replicate experiments at 18-, 4-, and 2-hour revealed that ProMag consistently achieved >90% labelling efficiency at the lowest concentration (20□µg Fe/mL), regardless of incubation duration. In contrast, VivoTrax required □240□µg Fe/mL to achieve similar efficiency. SynoMag-D and Ferumoxytol showed no detectable labelling without TAs (PS and heparin). SPION uptake is influenced by physicochemical properties, particularly surface coatings that modulate colloidal stability and cellular[19, 20]. ProMag, with its carboxyl-functionalized polystyrene coating and large hydrodynamic diameter (∼1□µm), may benefit from enhanced sedimentation and surface contact, facilitating endocytosis[21]. Conversely, SynoMag-D and Ferumoxytol labelling improved only in the presence of TAs after 18 hours. PS, a cationic TA, reverses the negative charge of SPIONs, promoting electrostatic interaction with the cell membrane and forming iron–PS nanocomplexes[12], which visible under light microscopy (Figure 1c-d). These aggregates may adhere to the cell surface or remain extracellular, with internalization likely during extended incubation. However, the increased hydrodynamic size and aggregation may impair MPI signal due to restricted Brownian motion, as previously reported in cancer cells[12, 22]. Consequently, SynoMag-D and Ferumoxytol were excluded from further MPI analysis.

Cell viability assays revealed a concentration-dependent decline for ProMag and VivoTrax, maintaining >80% and >90% viability at 20 and 120□µg Fe/mL (2-hour incubation), respectively. This highlights the importance of balancing efficient labelling with cytocompatibility to preserve MSC function. SynoMag-D and Ferumoxytol labelling showed variable viability, likely due to TA-induced membrane disruption[23, 24]. Notably, in iron-free control conditions, unbound TAs could interact more freely with cell membranes, possibly increasing cytotoxicity.

MPI characterization confirmed that there was a linear correlation between the number of labelled cells and total signal intensity for both ProMag and VivoTrax as it was shown in previous studies[8, 22]. Cells labelled with ProMag produced greater signal intensity than ones with VivoTrax, likely due to their higher iron uptake (∼28 pg/cell vs. ∼5.75 pg/cell). However, when total signal intensity was normalized to iron content, VivoTrax exhibited higher magnetization per unit of internalized iron compared to ProMag. This might be ascribed to physicochemical characteristics of the iron core and its MPI field response, which can be influenced by aggregation state, compartmentalization, and intracellular viscosity, all of which affect Néel and Brownian relaxation mechanisms[25]. While the detection sensitivity was noticeably higher for ProMag-labelled cells compared to those labelled with VivoTrax, the resolution, defined by the FWHM of the signal, remained comparable between the two. This suggests that ProMag labelling provides enhanced sensitivity without substantially compromising spatial resolution. Nevertheless, signal per unit iron favours VivoTrax, making it potentially more suitable in contexts where iron overload or cytotoxicity is a concern. The minimum cell detection limits for the MSCs labelled with ProMag and VivoTrax were determined as 12,500 and 50,000 cells, respectively. In another study, detection limits of 4T1 cancer cells labelled with VivoTrax was measured as 8000 cells with 2D MPI analysis[26]. This variation could be related to phagocytic activity and intracellular fate of the internalized nanoparticles and indicates the labelling efficiencies of the cells varies greatly depending on cell type.

In most studies it was shown that SPIONs were generally internalized through endocytosis, which is a temperature-sensitive process and known to be halted at temperatures lower than 10°C[19, 27]. Our low-temperature endocytosis inhibition assays confirmed the active, temperature-sensitive internalization pathway for both ProMag and VivoTrax. Furthermore, TEM evaluation of labelled cells showed that both particles were mostly localized in endosomes, reinforcing the interpretation that our labelling strategy is achieved by active transport rather than passive absorption. However, our TEM results revealed substantial differences in SPION localization and aggregation behaviour within the cells. ProMag particles were primarily found individually within endosomes, often in proximity but without forming aggregates, while VivoTrax particles exhibited a higher degree of aggregation within endocytic vesicles and appeared in various intracellular compartments, including mitochondria and endoplasmic reticulum. The intracellular distribution pattern may impact MPI signal performance: larger intracellular aggregates, as seen with VivoTrax, may restrict rotational freedom, leading to signal attenuation. In line with this, several studies have shown that the MPI signal tends to decrease after internalization, despite strong signals from free SPIONs[12, 21, 25, 28]. This highlights the importance of considering not only uptake but also intracellular behaviour when selecting SPIONs for MPI applications

The commercial SPIONs compared in this study have been used for MPI tracking in previous studies[21, 29]; however, they were originally developed for other purposes, such as MRI contrast agents or as medicinal supplements like Ferumoxytol, rather than for MPI, which operates based on different physical principles. Therefore, this comparison of SPIONs provides a valuable starting point for designing new SPIONs that can fully exploit the potential of MPI in future studies. In addition, special attention is required for their *in vivo* application, which presents unique challenges such as signal reduction after administration due to cell dispersion, positional variability, or changes in the tissue environment[21]. Moreover, *in vivo* imaging performance may not fully reflect *in vitro* results due to differences in aggregation, compartmentalization, or degradation pathways. Thus, the lack of *in vivo* analysis can be considered one of the limitations of our study. Furthermore, the removal of unbound SPIONs after labelling is critical. Larger particles, such as ProMag, tend to adhere more strongly to the cell membrane than do smaller particles, which may result in their detachment post-transplantation and uptake by host tissue cells. Therefore, an effective method for washing off unbound particles must also be carefully considered.

Realising the potential of MPI requires the optimization of SPION labelling strategies. Our results demonstrated that ProMag exhibited the highest labelling efficiency, iron uptake, and MPI signal intensity of the tested SPIONs, positioning it as a strong candidate for future cell tracking applications. However, VivoTrax produced a superior signal per unit of iron, suggesting that enhancing its cellular uptake could lead to even better signal generation. Our findings underscore that both uptake efficiency and the intracellular fate of SPIONs critically influence MPI performance.

## Statement & Declarations

### Author contributions

Serbay Ozkan and Patricia Murray contributed to the study conception and design. Material preparation, data collection and analysis were performed by Serbay Ozkan, Elena Ureña Horno, Mahon L Maguire and Reilteann Saul. The first draft of the manuscript was written by Serbay Ozkan and all authors commented on previous versions of the manuscript. All authors read and approved the final manuscript.

### Data Sharing Statement

The data that support the findings of this study are available from the corresponding author upon reasonable request.

### Artificial intelligence use statement

No artificial intelligence (AI) or AI-assisted technologies were used in the preparation of this article.

