## Supplementary Information for "Evaluating Labelling Efficiency of Commercial SPIONs in Mesenchymal Stem/Stromal Cells for Magnetic Particle Imaging Applications"

**Running title**: Comparison of SPIONs for magnetic particle imaging

^1^ Women’s and Children’s Health Department, Faculty of Health and Life Sciences, University of Liverpool, Liverpool L69 3GE, UK

^2^ Histology and Embryology Department, Faculty of Medicine, Izmir Kâtip Celebi University, Izmir Turkey

^3^ Department of Chemistry, University of Liverpool, Liverpool L69 7ZD, UK

^4^ Centre for Preclinical Imaging, University of Liverpool, Liverpool L69 3BX, UK

^5^ Department of Molecular and Clinical Cancer Medicine, Faculty of Health and Life Sciences, University of Liverpool, Liverpool L69 3BX, UK

**Corresponding author:** Serbay Ozkan (PhD)

**Adress:** Women’s and Children’s Health Department, Faculty of Health and Life Sciences, University of Liverpool, Liverpool L69 3GE, UK.

**Supplementary Figures**

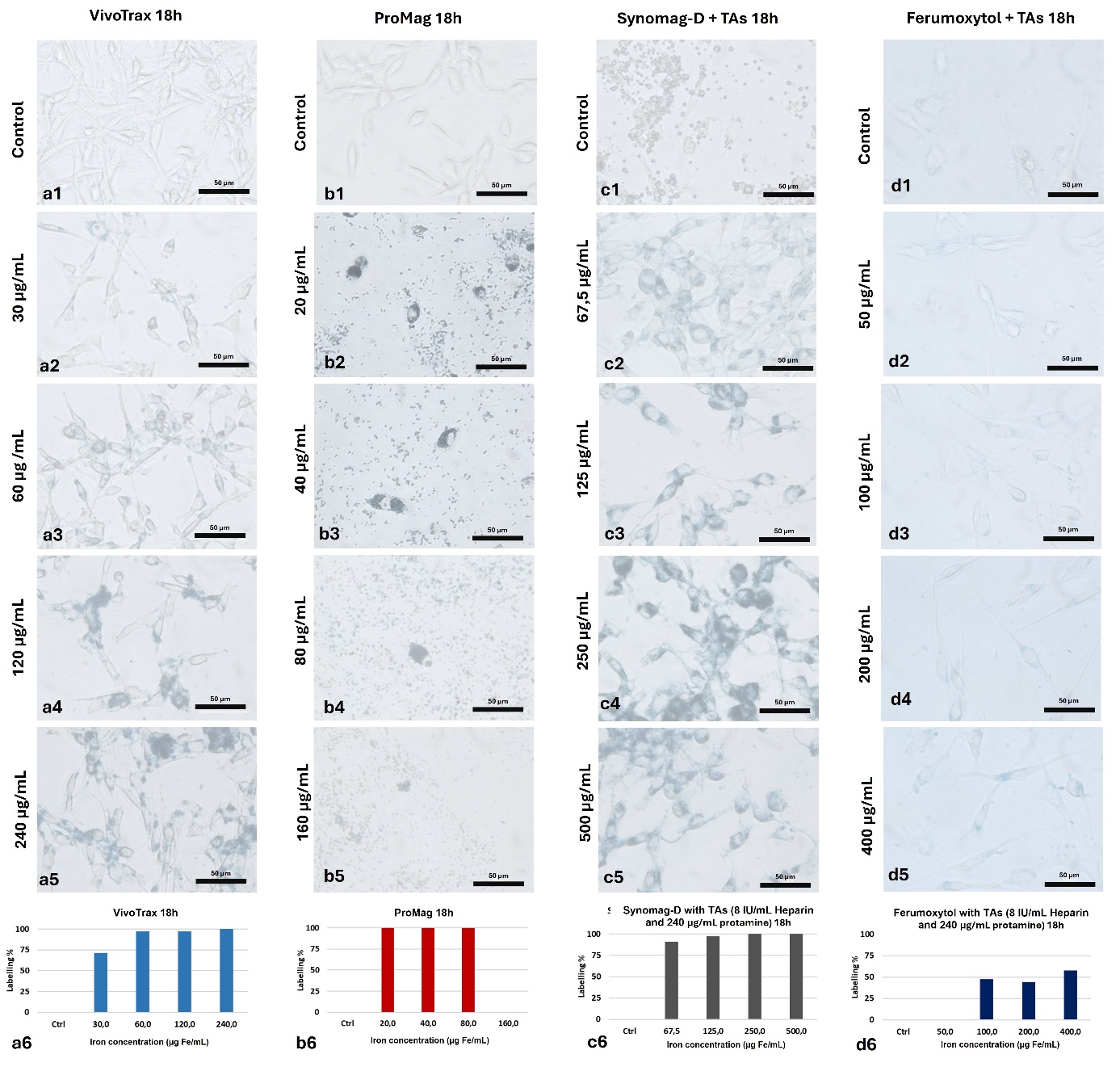

**Supplementary Figure 1**: (a-d) Representative figures of adherent mBM-MSCs incubated with different commercial magnetic particle for 18-hours at different ranges of concentrations and graphical expression of labelling efficiencies (a6, b6, c6, and d6)

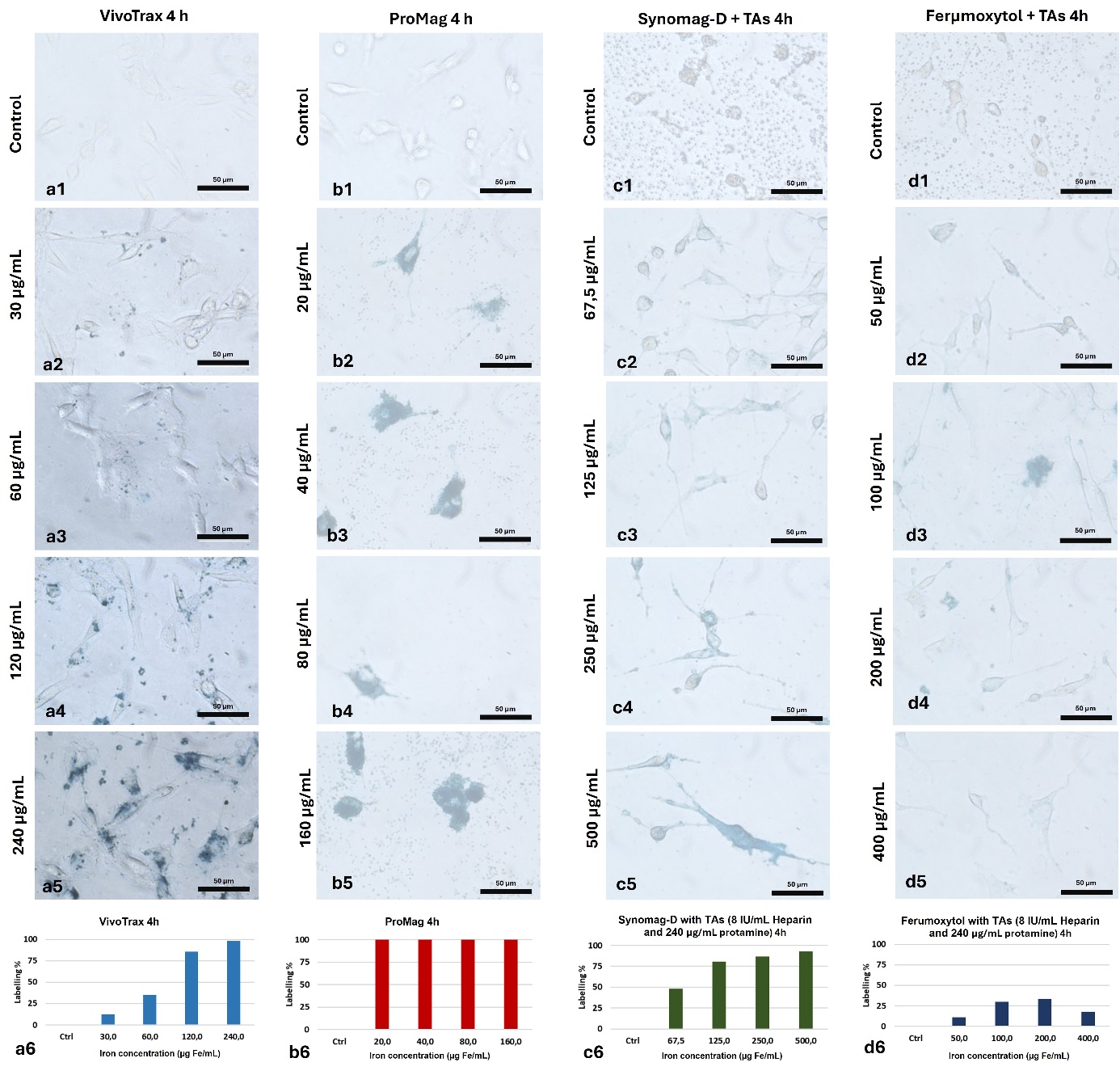

**Supplementary Figure 2**: (a-d) Representative figures of adherent mBM-MSCs incubated with different commercial magnetic particle for 4-hours at different ranges of concentrations and graphical expression of labelling efficiencies (a6, b6, c6, and d6)

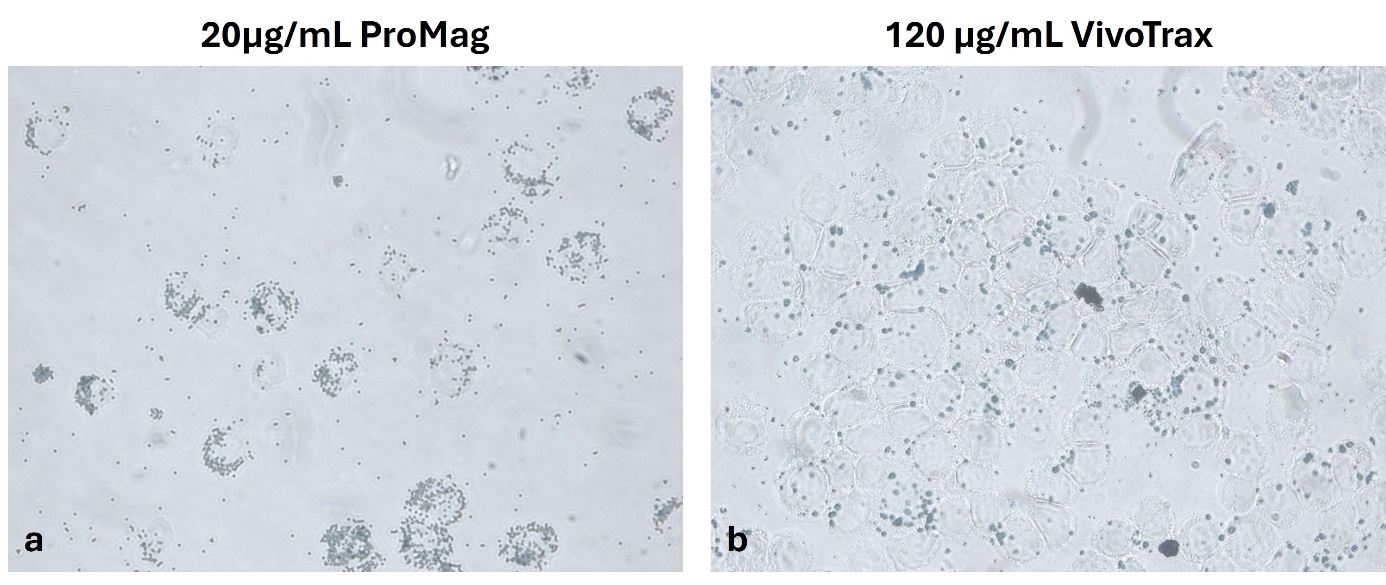

**Supplementary Figure-3:** (a-b) Demonstrative pictures for ProMag (a) and VivoTrax (b) labelling prior to MPI analysis.

**Supplementary Methods**

**Supplementary method-1**: Specifically, a line profile was drawn through the image along the axis of maximum signal intensity, using MagImage Image Analysis Software (Magnetic Insight). The resulting profile was fitted to a Lorentzian curve, and the Full Width at Half Maximum was extracted to quantify the resolution in millimetres.

**Supplementary method-2**: Following secondary fixation with 1% osmium tetroxide (OsO_4_), samples were treated with potassium ferrocyanide for 1 hour at RT. The cells were then rinsed six times with double-distilled water (ddH_2_O), for five minutes each rinse. After overnight incubation with 0.75% uranyl acetate in ddH_2_O at 4 °C, the cells were rinsed three times. Subsequently, the cells were dehydrated using a graded ethanol series (30%, 50%, 70%, 90%, 100%) for 10 minutes each. They were then treated with resin-ethanol mixtures (1:2 and 2:1 resin/ethanol), followed by three incubations in 100% resin, and embedded in resin by overnight incubation at 60 °C. Semi-thin sections were cut to identify regions of interest for ultra-thin sectioning, after which 40–50 nm thick sections were collected on copper grids. The grids were contrasted with 2% uranyl acetate for 3–5 minutes, followed by lead citrate for 3–5 minutes.
